## Supplementary material for "The association between *Dioscorea sansibarensis* and *Orrella dioscoreae* as a model for hereditary leaf symbiosis": Figure S1

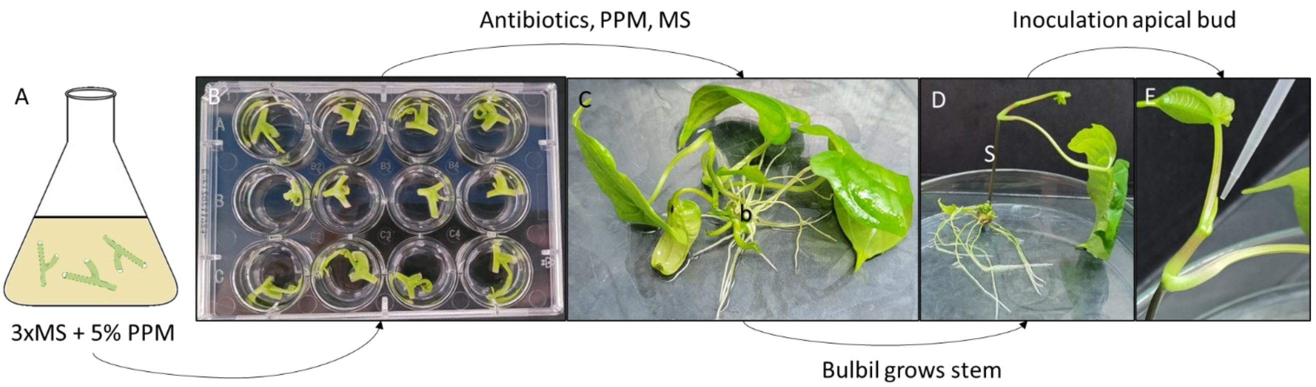


**Figure S1: Method developed to make aposymbiotic plants and re-introduce a bacterium of interest. (A)** Node cuttings were taken from adult plants and incubated for 8 hours in 5% PPM for initial sterilization. **(B)** Node cuttings were incubated in a mixture of liquid MS, antibiotics and PPM for 3 weeks. **(C)** After 3-4 weeks, a bulbil (b) with its root system became apparent. Multiple leaves have formed from the node and are providing sugars to the plant. **(D)** The bulbil grows its own stem (S) that uses gravitropism to grow up and after the emergence of two leaves, the apical bud becomes visible. **(E)** After confirmation of being aposymbiotic by crushing and plating out the newly developed acumen(s), the plant was re-inoculated with a bacterium of interest by dropping 2 µl of the bacterial suspension on the apical bud.
