## Supplementary material for "The association between *Dioscorea sansibarensis* and *Orrella dioscoreae* as a model for hereditary leaf symbiosis": Figure S2

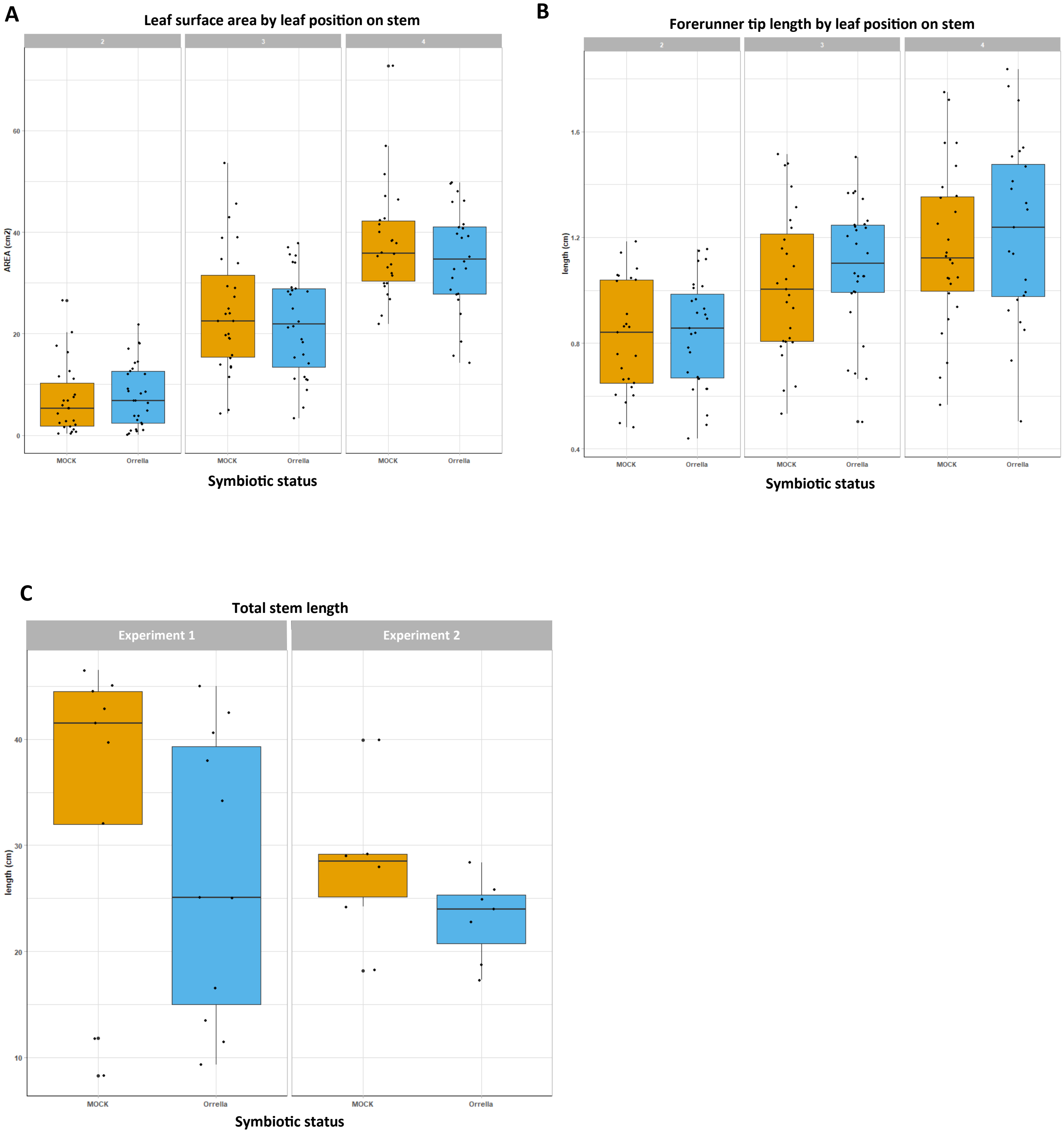


**Figure S2: Morphological parameters of aposymbiotic vs. symbiotic *D. sansibarensis* in gnotobiotic conditions.** Wild-type colonized *D. sansibarensis* were inoculated by a *O. dioscoreae* R-71412 cell suspension (Orrella) or a sterile 0.4% NaCl solution (MOCK) and grown for 4 weeks in gnotobiotic conditions. Leaf surface area (A) and length of the forerunner tip containing the bacterial glands (B) were measured for 3 leaves per plant, starting with the leaf closest to the shoot tip (leaf 1, not shown). C. Total stem length measured from the crown to the shoot tip. Data from 2 independent experiments are shown separately. Data from mock-inoculated plants are shown in orange, and in blue for *O. dioscoreae*-inoculated plants. The distributions of values between the *O. dioscoreae*– or mock-inoculated plants are identical for each of the 3 parameters (Wilcoxon rank sum test $p >$ 0.05).
