## Supplementary material for "The association between *Dioscorea sansibarensis* and *Orrella dioscoreae* as a model for hereditary leaf symbiosis": Figure S3

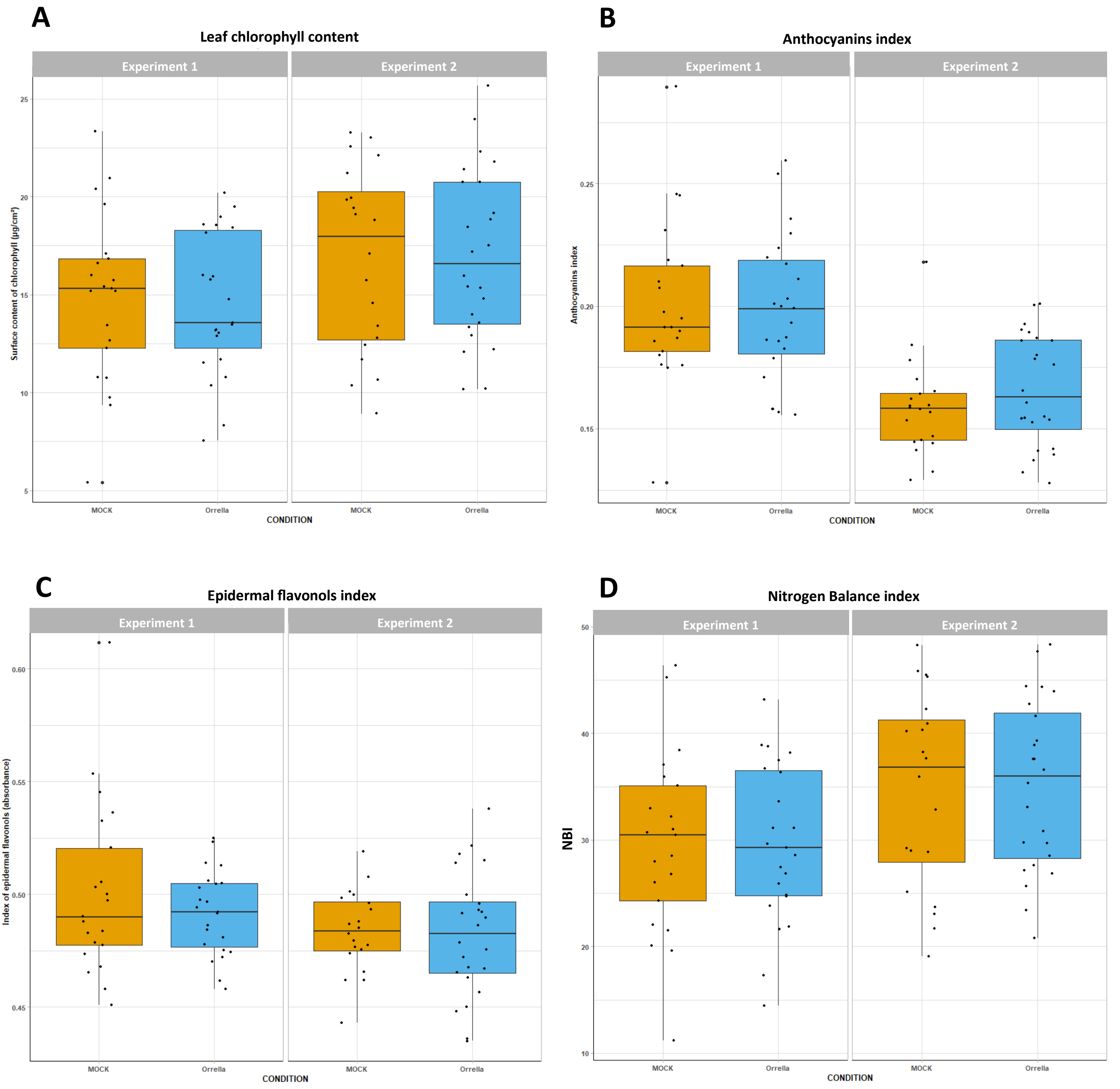


**Figure S3: Physiological parameters of aposymbiotic vs. symbiotic *D. sansibarensis* in gnotobiotic conditions.** Wild-type colonized *D. sansibarensis* were inoculated by a *O. dioscoreae* R-71412 cell suspension (Orrella) or a sterile 0.4% NaCl solution (MOCK). Physiological parameters were measured using a hand-held optical meter after 4 weeks of growth in gnotobiotic conditions. Parameters measured include **A**. Chlorophyl content (Chl); **B**. Anthocyanins index, measured as a function of green light absorbed by the sample; **C**. Flavonoids index (Flav), measured as a function of UV light absorbed by the sample and **D**. Nitrogen Balance Index (NBI) is measured as the ratio of Chl and Flav and is an indicator of C/N allocation changes due to N-deficiency. Data from 2 independent experiments are shown separately. Data from mock-inoculated plants are shown in orange, and in blue for *O. dioscoreae*-inoculated plants. The distributions of values between the *O. dioscoreae*– or mock-inoculated plants are identical for each of the 4 parameters (Wilcoxon rank sum test $p >$ 0.05).
