## Supplementary material for "The association between *Dioscorea sansibarensis* and *Orrella dioscoreae* as a model for hereditary leaf symbiosis": Table S1

**Table S1: Bacterial species used in this study**

| Species | Strain or plasmid | Description | Growth conditions | Reference or source |
|---|---|---|---|---|
| *Orrella dioscoreae* | R-71417 | Strain R71412 with mini Tn7(Gm) Ptac-mCherry | TSA + Nalidixic acid 30 µg/ml + Gentamy-cin 20 µg/ml, 28°C, aerobic | This study |
| *Orrella dioscoreae* | R-71416 | Strain R71412 with mini Tn7(Gm) Ptac-GFP | TSA + Nalidixic acid 30 µg/ml + Gentamy-cin 20 µg/ml, 28°C, aerobic | This study |
| *Orrella dioscoreae* | R-67584 | Accession nr: 19760001, plant from direct wild origin, Origin: Congo DR | TSA, 28°C, aerobic | Evrard C., Lou-vain-la-Neuve, U.C.L., Ecologie et Biogeographie, |
| *Orrella dioscoreae* | R-67173 | Isolated from leaf nodules of *Di-oscorea sansibarensis* (Zanzibar yam) | TSA, 28°C, aerobic | This study |
| *Orrella dioscoreae* | R-67088 | isolate from leaf nodules of *Di-oscorea sansibarensis,* botanical garden of the University of Ghent, Gent, Belgium | TSA, 28°C, aerobic | This study |
| *Orrella dioscoreae* | R-67090 | isolate from leaf nodules of *Di-oscorea sansibarensis,* botanical garden of the University of Ghent, Gent, Belgium | TSA, 28°C, aerobic | This study |
| *Orrella dioscoreae* | LMG-29303$^T$ | Type strain *O. dioscoreae* | TSA, 28°C, aerobic | This study |
| *Escherichia coli* | Top10 | | LB, 37°C | |
