## Supplementary material for "The association between *Dioscorea sansibarensis* and *Orrella dioscoreae* as a model for hereditary leaf symbiosis": Table S2

**Table S2: Minimum inhibitory concentrations of biocidal products on different *O. dioscoreae* strains**

| | Carbenicillin | Cefotaxime | PPM |
|---|---|---|---|
| **R-67173** | 64 | 128 | 0,04 |
| **R-67584** | 64 | 128 | 0,04 |
| **R-67088** | 32 | 128 | 0,04 |
| **R-67090** | 32 | 128 | 0,04 |
| **LMG 29303$^T$** | 32 | 128 | 0,04 |
