## Supplementary material for "The association between *Dioscorea sansibarensis* and *Orrella dioscoreae* as a model for hereditary leaf symbiosis": Table S3

**Table S3: Symbiotic status of plants used in phenotyping experiment.** APO= aposymbiotic status, SYM= symbiotic status, check-ups quantified the amount *of O. dioscoreae* found in new leaf acumens. Not= Majority isolates not identified as *O. dioscoreae.* Last column gives the eventual identity given to the sample for further analysis: APO= aposymbiotic plant, unknown= colonized by bacteria other than *O. dioscoreae, Orrella dioscoreae* = colonized by *Orrella dioscoreae*

| Plant ID | Symbiotic status | inoculated with | Status T0 | Status T1 | Status T2 | check-up 3B | Status 3 Identification | Final status |
|---|---|---|---|---|---|---|---|---|
| 1 | APO | Mock | APO | 0 | 0 | 0 | | APO |
| 2 | APO | *O. dioscoreae* | APO | 0 | 0 | 0 | | APO |
| 4 | SYM | | SYM | | 0 | 0 | | Unknown |
| 5 | APO | Mock | APO | 0 | 0 | 0 | | APO |
| 6 | APO | Mock | APO | 0 | 0 | 0 | | APO |
| 8 | SYM | | | | 0 | 0 | | Unknown |
| 9 | APO | *O. dioscoreae* | APO | 0 | $10^4$ | $10^5$ | not | Unknown |
| 10 | SYM | | APO | | 0 | $10^2$ | not | Unknown |
| 11 | APO | Mock | APO | 0 | 0 | 1 | not | APO |
| 12 | APO | *O. dioscoreae* | APO | 0 | $10^3$ | $10^3$ | *O. dioscoreae* | *O. dioscoreae* |
| 13 | APO | Mock | APO | 0 | 0 | $10^2$ | not | APO |
| 14 | APO | *O. dioscoreae* | APO | 0 | 0 | $10^2$ | *O. dioscoreae* + other | Unknown |
| 15 | APO | Mock | sym | 0 | $10^3$ | $10^3$ | not | Unknown |
| 16 | APO | *O. dioscoreae* | O. dioscoreae | $10^3$ | $10^3$ | $10^3$ | *O. dioscoreae* | *O. dioscoreae* |
| 17 | APO | *O. dioscoreae* | *O. dioscoreae* | 0 | $10^3$ | $10^3$ | *O. dioscoreae* | *O. dioscoreae* |
| 19 | SYM | | SYM | | 0 | $10^2$ | *O. dioscoreae* | *O. dioscoreae* |
| 20 | APO | Mock | SYM | 0 | 0 | $10^1$ | *O. dioscoreae* + other | Unknown |
| 21 | SYM | | SYM | | 0 | $10^2$ | *O. dioscoreae* + other | *Orrella dioscoreae* |
| 22 | APO | *O. dioscoreae* | indecisive | 0 | 1 | $10^2$ | *O. dioscoreae* + other | *O. dioscoreae* |
| 23 | SYM | | SYM | | 0 | 0 | | Unknown |
| 24 | SYM | | SYM | | 0 | 0 | | Unknown |
